# 4-Phenylbutyric Acid Activates an NF-κB - Egr-1 Axis to Control Myoblast Proliferation and ECM Gene Expression Profiles

**DOI:** 10.64898/2026.01.18.700214

**Authors:** Kana Tominaga, Naoomi Tominaga

**Affiliations:** Division of Neuropsychiatry, Department of Neuroscience, Yamaguchi University Graduate School of Medicine, 1-1-1 Minami-kogushi, Ube, Yamaguchi, 755-8505, Japan; Division of Clinical Laboratory Sciences, Department of Nursing and Laboratory Science, Yamaguchi University Graduate School of Medicine, 1-1-1 Minami-kogushi, Ube 755-8505, Japan

**Author notes:** Corresponding author: Kana Tominaga, Ph.D. Division of Neuropsychiatry, Department of Neuroscience, Yamaguchi University School of Medicine, 1-1-1 Minami-kogushi, Ube, Yamaguchi, 755-8505, Japan.

## Abstract

Myoblast proliferation and differentiation are tightly controlled by epigenetic mechanisms, yet how clinically used epigenetic modulators influence myogenic cell fate remains incompletely understood. Here, we demonstrate that the histone deacetylase inhibitor and chemical chaperone 4-phenylbutyric acid (4-PBA) selectively promotes myoblast proliferation without inducing differentiation in C2C12 cells. Mechanistically, 4-PBA increases histone H3 acetylation at lysines 18 and 27 via downregulation of HDAC5, resulting in activation of NF-κB p65. Chromatin immunoprecipitation identifies early growth response 1 (Egr-1) as a direct transcriptional target of NF-κB p65. Transcriptomic analyses reveal that Egr-1 regulates extracellular matrix- and myogenesis-associated gene programs, including multiple collagen genes. Consistently, 4-PBA induces a transcriptional signature that significantly overlaps with Egr-1-dependent gene expression. Functional studies further establish that the NF-κB p65 - Egr-1 axis is required for 4-PBA-mediated transcriptional remodeling in proliferating myoblasts. Together, these findings uncover an epigenetic mechanism by which 4-PBA modulates myoblast proliferation through HDAC5-dependent histone acetylation and NF-κB p65 - Egr-1 driven transcriptional programs, providing insight into how epigenetic therapeutics influence skeletal muscle cell behavior.

**Highlight:**

- 4-PBA enhances murine myoblast proliferation independently of differentiation induction.
- 4-PBA enhances H3K18 and H3K27 acetylation through downregulation of HDAC5.
- NF-κB p65 activates Egr-1 by directly binding to the Egr-1 promoter region.
- 4-PBA stimulates the HDAC5 - NF-κB p65 - Egr-1 axis drives extracellular matrix-related gene profiles.

## Introduction

Myoblasts are proliferative precursor cells derived from muscle satellite cells^1,2^ Satellite cells are skeletal muscle stem cells that reside in a quiescent state beneath the basal lamina of muscle fibers^3,4^. In response to muscle injury or growth signals, satellite cells become activated and give rise to myoblasts, which proliferate and subsequently differentiate to form new muscle fibers (myotubes/myofibers)^5,6^. This process is essential for skeletal muscle growth, regeneration, and repair and represents a promising therapeutic target for muscle-related diseases^7,8^.

Histone acetylation plays a central role in the proliferation and the differentiation of myoblasts^9^. During differentiation, histone acetyltransferases (HATs) are recruited to the promoters and enhancers of muscle-specific genes, leading to increased acetylation of histone marks such as H3K9ac and H3K27ac^10–12^. This epigenetic remodeling activates key myogenic genes, including Myog, myosin heavy chain (Myh), and muscle creatine kinase^10,13^. The myogenic transcription factor MyoD recruits HATs such as p300/CBP and PCAF to muscle gene loci, enhancing chromatin accessibility and reinforcing muscle-specific transcription^14,15^. Concurrently, histone deacetylases (HDAC) are displaced from muscle gene promoters but remain associated with cell cycle-related genes to ensure permanent withdrawal from proliferation^16,17^. Pharmacological inhibition of HDAC increases histone acetylation and can accelerate myoblast differentiation, underscoring the importance of balanced acetylation in myogenesis^18^.

4-PBA is currently used clinically to treat patients with urea cycle disorders under the names Buphenyl (sodium phenylbutyrate) and Ravicti (glycerol phenylbutyrate)^19^. Its primary mechanism of action involves binding toxic ammonia to provide an alternative pathway for its elimination^19,20^. In addition, 4-PBA functions as a chemical chaperone that reduces endoplasmic reticulum (ER) stress and prevents protein aggregation^21–23^. 4-PBA has attracted interest for disorders involving protein misfolding, such as cystic fibrosis with the ΔF508 *CFTR* mutation and vascular ehlers danlos syndrome with *COL3A1* mutations^24–26^. Previous studies have shown that 4-PBA treatment restores the membrane localization of pathogenic mutant dysferlin (Dysf) in cultured myotubes and myofibers obtained from dysferlinopathy model mice, suggesting that 4-PBA rescues dysferlin from aggregation or degradation in the endoplasmic reticulum (ER)^27^. 4-PBA is also recognized as a HDAC inhibitor, adding an epigenetic mechanism to its therapeutic effects. During cardiomyogenesis, 4-PBA promotes early cardiac differentiation via increased histone acetylation but inhibits late-stage differentiation by maintaining the expression of pluripotency genes such as *Oct4, Nanog,* and *Sox2*, demonstrating that HDAC regulation exerts context-dependent effects^28^. In the case of cancer, 4-PBA inhibits tumor cell proliferation by increasing histone acetylation, inducing epithelial-mesenchymal transition, and promoting acetyl-H3-mediated transcription^29–31^. Hence, these effects suggest that 4-PBA may alleviate ER stress caused by misfolded proteins and enhance protein homeostasis. In contrast, despite the clinical response to 4-PBA in diseases caused by misfolded proteins, little is known about its role as an HDAC inhibitor of myogenesis.

Our study demonstrated that 4-PBA regulates myoblast differentiation through epigenetic and transcriptional mechanisms. In C2C12 myoblasts, chromatin immunoprecipitation assays revealed that NF-κB p65 directly binds to the Egr-1 promoter, identifying Egr-1 as a transcriptional target of the NF-κB signaling pathway. During myogenesis, 4-PBA treatment markedly increased histone H3 acetylation, especially at Lys18 and Lys27, and downregulated HDAC5 expression, resulting in increased Egr-1 and p65 expression. Moreover, transcriptome analysis revealed that Egr-1 regulated extracellular matrix (ECM)-related genes, including multiple collagen genes. This study revealed that 4-PBA acts as an HDAC inhibitor that contributes to myogenic homeostasis by regulating the NF-κB - Egr-1 axis.

## Results

### 4-PBA induces cell proliferation in murine myoblasts

To confirm the effects of 4-PBA on myoblast differentiation, we first examined morphological changes in C2C12 myoblasts following 4-PBA treatment. C2C12 myoblasts were cultured under differentiation conditions and treated with 4-PBA for four days (Figure S1A). Myotube formation was then assessed by measuring the myofusion index and the protein levels of myosin heavy chain 1 (Myh1), a major contractile protein, using immunofluorescence analysis (Figure 1A). We observed a significant decrease in the fusion index and reduced Myh1 expression in 4-PBA-treated cells compared with control cells (Figure 1B). We next examined the protein expression of myogenin (Myog), a myogenic transcription factor, and Dysf, a marker of mature muscle fibers, under 4-PBA-treated conditions (Figure 1C). The results showed that 4-PBA significantly suppressed the induction of Myog at all stages (day 1 - 4) and Dysf at later stages (day 3 - 4) of myotube differentiation (Figure 1D). On the other hand, RNA-seq analysis of myoblast differentiation from day 0 to day 4 indicated that there was no difference in the expression of myoblast differentiation-related genes between DMSO- and 4-PBA-treated cells (Figure S1B - S1D). These results suggest that 4-PBA suppresses myogenic differentiation primarily through post-transcriptional, translational, or protein stability-related mechanisms rather than by regulating myogenic gene expression at the mRNA level.

**Figure 1.**
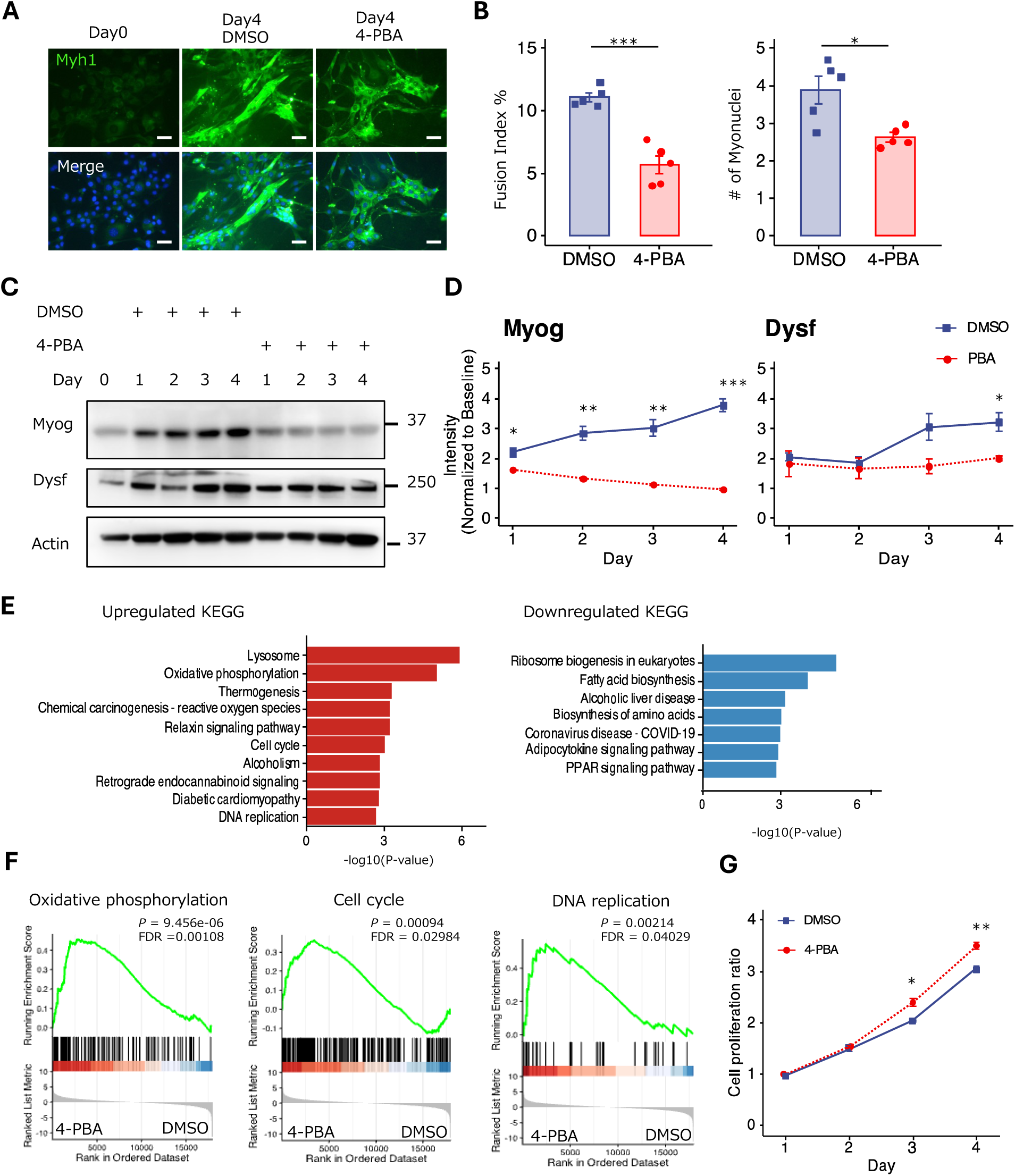
4-PBA Promotes Myoblast Proliferation but Impairs Myogenic Differentiation. (A) Representative immunofluorescence images of C2C12 myotubes stained for Myh1 (green) and nuclei (blue). Scale bar, 400 µm. (B) (left) Quantification of Myh1⁺ myotube area per field 4 days after differentiation (n = 5). (right) the number of nuclei within multinucleated myotubes (n = 5). (C) Representative immunoblots showing Myogenin (Myog) and Dysferlin (Dysf) expression in C2C12 cells treated with 1 mM 4-PBA or DMSO for the indicated time points. (D) Quantification of Myog and Dysf protein levels shown in (C), normalized to β-actin (Actin). Data are representative of three independent experiments. (E) KEGG pathway enrichment analysis of differentially expressed genes in C2C12 myotubes at day 4 after differentiation, comparing treatment with 1 mM 4-PBA and DMSO. (F) Gene set enrichment analysis (GSEA) plots for the indicated KEGG pathways in 4-PBA– or DMSO-treated cells. FDR, false discovery rate. (G) Proliferation of C2C12 myoblasts treated with 1 mM 4-PBA or DMSO. Data in (B), (D) and (G) are presented as mean ± S.E.M. and analyzed using an unpaired two-tailed Student’s t test. \**P* < 0.05, \*\**P* < 0.01, \*\*\**P* < 0.001.

To identify the biological pathways affected by 4-PBA treatment in myoblasts, we analyzed RNA-seq data (GSE315556) from C2C12 myoblasts treated with 4-PBA under differentiation conditions for four days. KEGG pathway analysis revealed that genes upregulated by 4-PBA treatment were primarily associated with cell proliferation-related pathways (Figure 1E and Data S1). Three representative pathways were visualized to facilitate interpretation (Figure 1F). To validate the RNA-seq results indicating enhanced proliferation, we examined cell proliferation in C2C12 myoblasts treated with 4-PBA or DMSO as a control. Compared with DMSO-treated cells, C2C12 myoblasts treated with 4-PBA showed significantly increased proliferation after four days (Figure 1G), suggesting that compared with myoblast differentiation, 4-PBA promotes cell proliferation.

### 4-PBA induces Egr-1 expression during myogenesis

4-PBA acts as an HDAC inhibitor, thereby increasing histone acetylation and consesquently promoting gene expression. Therefore, to identify genes upregulated by 4-PBA treatment during myogenic differentiation, we compared the gene expression profiles of 4-PBA-treated C2C12 myoblasts with those of DMSO-treated or undifferentiated (baseline) cells via RNA-Seq (GSE315556, Data S2). Principal component analysis (PCA) revealed that compared with the baseline treatment, the 4-PBA and DMSO treatments caused significant changes in the state, after which the 4-PBA-treated C2C12 myotubes formed a distinct cluster that was clearly separated from the DMSO treatment (Figure 2A). These results indicated a strong and consistent global transcriptional change caused by 4-PBA treatment during myogenesis. Transcriptomic analysis revealed that only two genes, Egr-1 and Fos, were differentially expressed genes (DEGs) based on the criteria of | log2-fold change | > 1.2 and *p* < 0.01 (Figure 2B and 2C). The mRNA expression levels of both genes were greater in 4-PBA-treated C2C12 myotubes than in DMSO-treated cells or baseline controls (Figure 2D). At the protein level, Egr-1 expression markedly increased in 4-PBA-treated C2C12 cells compared with that in DMSO-treated and control cells, particularly during the early stages of differentiation (Figure 2E and 2F). In contrast, Fos protein expression did not significantly differ between 4-PBA- and DMSO-treated C2C12 cells. These findings indicated that Egr-1 is a primary and functionally relevant target of 4-PBA in the transformation of C2C12 myoblasts to myotubes.

**Figure 2.**
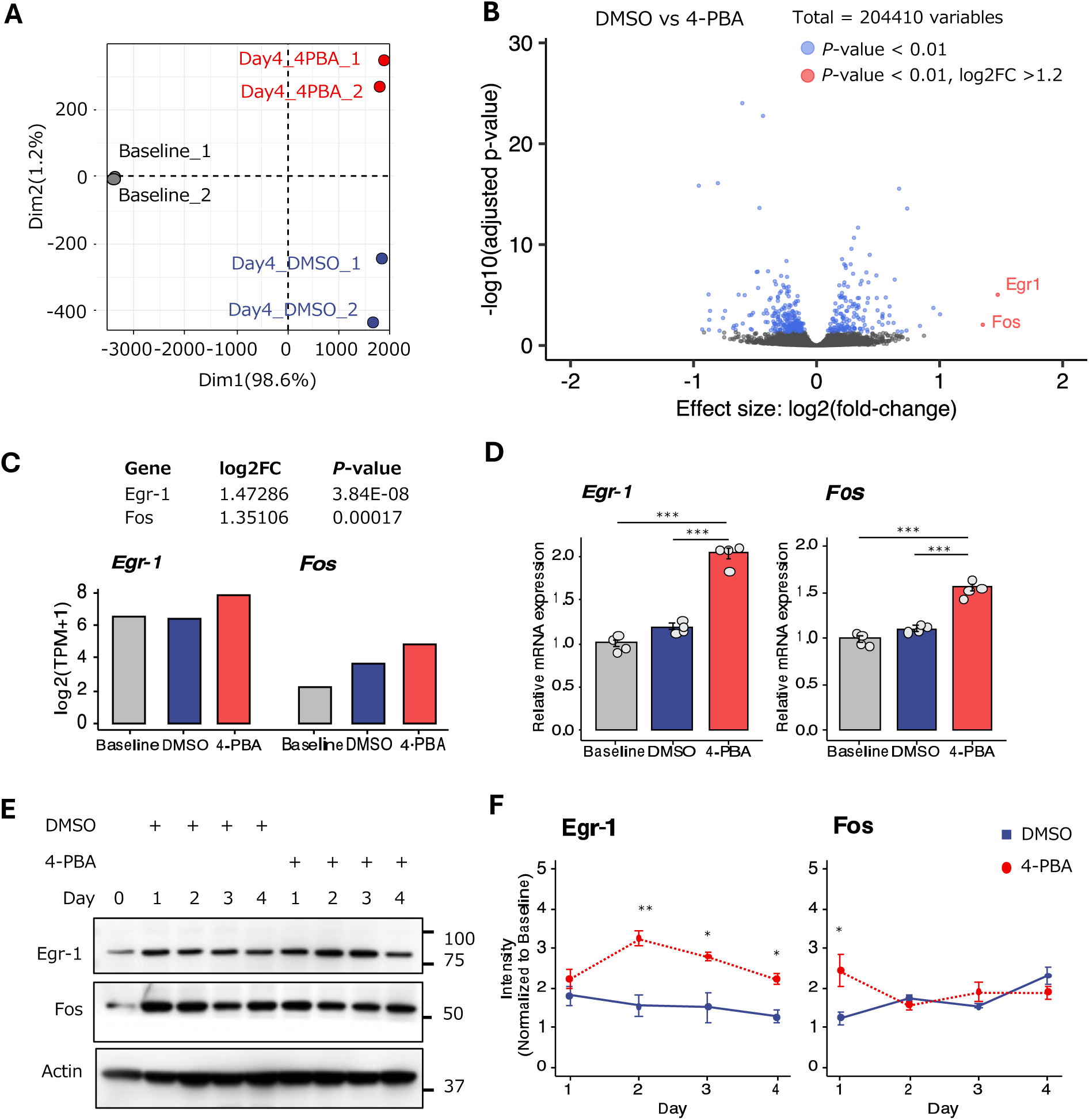
4-PBA Induces Egr-1 Expression during Myoblast Differentiation. (A) Principal component analysis (PCA) showing sample clustering based on differentially expressed genes in C2C12 cells treated with 1 mM 4-PBA or DMSO. Baseline indicates cells prior to differentiation induction. (B) Volcano plot of differentially expressed genes identified by RNA-seq analysis comparing 4-PBA–treated and DMSO-treated C2C12 cells following myotube differentiation after 4 days. Significantly upregulated genes are shown in red. FC, fold change; P-adj, adjusted p value. (C) RNA-seq–based expression levels of *Egr-1* and *Fos* mRNA in C2C12 cells treated with 1 mM 4-PBA or DMSO, shown as transcripts per million (TPM). (D) Quantitative PCR (qPCR) validation of *Egr-1* and *Fos* mRNA expression in C2C12 cells treated with 1 mM 4-PBA or DMSO. Data represent four independent experiments. (E) Representative immunoblots showing Egr-1 and Fos protein expression in C2C12 cells treated with 1 mM 4-PBA or DMSO for the indicated time points. (F) Quantification of protein levels shown in (E), normalized to β-actin (Actin). Data represent three independent experiments. Data in (D) and (F) are presented as mean ± S.E.M. and analyzed using one-way ANOVA with Tukey’s post hoc test (D) and unpaired two-tailed Student’s t test (F). \**P* < 0.05, \*\**P* < 0.01, \*\*\**P* < 0.001.

### 4-PBA modulates the NF-κB p65-dependent expression of murine myoblasts

To identify the signaling pathway that regulates Egr-1 expression, we focused on the Erk pathway and the NF-κB p65 signaling pathway. Some studies have shown that 4-PBA inhibits the activation of both pathways in cells, primarily through its ability to reduce ER stress. We treated C2C12 myoblasts with 1 mM 4-PBA or DMSO for four days during myotube differentiation and then evaluated the phosphorylation of Erk or NF-κB p65. Erk1/2 phosphorylation remained unchanged following 4-PBA or DMSO treatment, whereas NF-κB p65 expression and phosphorylation increased in 4-PBA-treated C2C12 cells from the early stages of differentiation (Figure 3A and 3B). To confirm whether Egr-1 expression is regulated by the canonical NF-κB signaling pathway, we analyzed the protein levels of Egr-1 in 4-PBA-treated C2C12 myoblasts treated with dehydroxymethylepoxyquinomicin (DHMEQ), an NF-κB p65 inhibitor, for 24 hours. Immunoblotting revealed that Egr-1 protein levels increased in response to 4-PBA treatment but decreased in response to the addition of DHMEQ (Figure 3C and 3D). NF-κB p65 may act as a transcription factor that regulates Egr-1.

**Figure 3.**
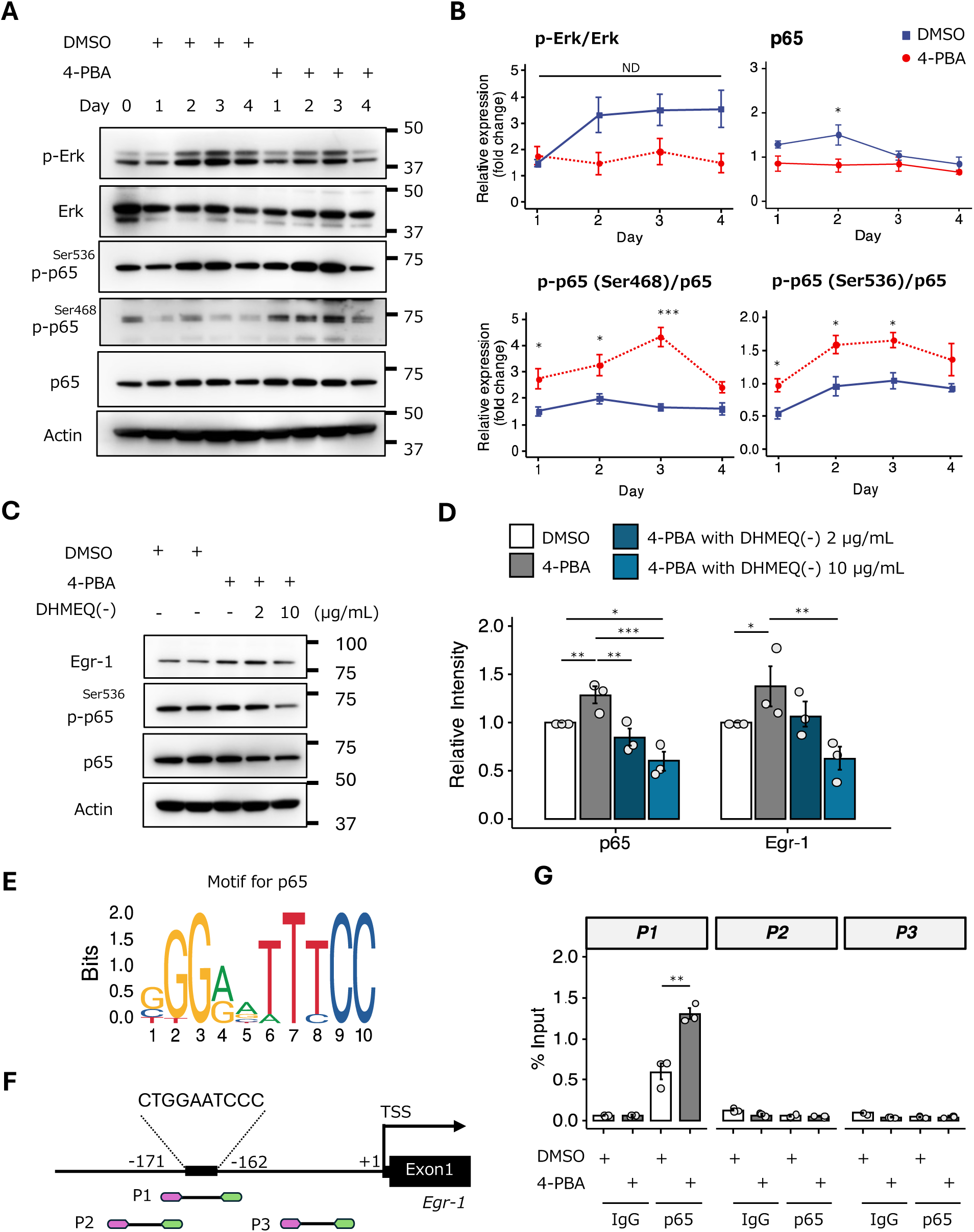
4-PBA Regulates NF-κB–Dependent Egr-1 Expression in C2C12 Cells. (A) Representative immunoblots showing total Erk1/2, phosphorylated Erk1/2 (Thr202/Tyr204), total p65, and phosphorylated p65 (Ser468 and Ser536) in C2C12 cells treated with 1 mM 4-PBA or DMSO for the indicated time points. β-actin (Actin) was used as a loading control. (B) Quantification of phosphorylated Erk1/2 (Thr202/Tyr204) and phosphorylated p65 (Ser468 and Ser536) shown in (A). Band intensities were normalized to total Erk1/2 or p65, respectively. Data represent three independent experiments. (C) Representative immunoblots showing Egr-1 and p65 protein levels in C2C12 cells treated with 1 mM 4-PBA in the presence or absence of the NF-κB inhibitor DHMEQ. Actin was used as a loading control. (D) Quantification of Egr-1 and p65 protein levels shown in (C), normalized to Actin. Data represent three independent experiments. (E) NF-κB p65 consensus binding motif predicted using the JASPAR database. (F) Predicted p65 binding sites within the Egr-1 promoter region identified by JASPAR analysis. (G) Chromatin immunoprecipitation (ChIP) analysis showing enrichment of the Egr-1 promoter regions (P1 and P2) in p65- or IgG-immunoprecipitated chromatin from 4-PBA–treated C2C12 cells. Data in (B), (D) and (G) are presented as mean ± S.E.M. and analyzed using an unpaired two-tailed Student’s t test (B) and one-way ANOVA with Tukey’s post hoc test (D, G). \**P* < 0.05, \*\**P* < 0.01, \*\*\**P* < 0.001; ns, not significant.

Putative NF-κB (p65) binding sites were identified based on the established κB consensus sequence (5′-GGGRNNYYCC-3′) in the *Homo sapiens* (GRCh38). However, within the EGR promoter region, a consensus NF-κB (p65) binding motif was identified in the human sequence, whereas no corresponding motif was detected in the mouse sequence. To identify NF-κB p65 binding sites that control Egr-1 transcription in the *Mus musculus* genome (GRCm39), we predicted putative NF-κB p65 binding sites within the −2000 bp promoter region of *Egr-1* using the JASPAR database (Figure 3E and S2A). We designed two primer pairs encompassing the predicted motifs (P1) and two primer pairs flanking regions adjacent to the motifs (P2 and P3) to assess DNA enrichment (Figure 3F and S2B). ChIP‒qPCR analysis revealed that the P1 fragment was directly bound by NF-κB p65, whereas the other fragments (P2 and P3) were not enriched (Figure 3G). These results demonstrate that NF-κB p65 directly binds to the *Egr-1* promoter, specifically at the P1 region (CTGGAATCCC), indicating that *Egr-1* is a direct transcriptional target of NF-κB p65. Moreover, compared with DMSO-treated C2C12 myoblasts, 4-PBA-treated C2C12 myoblasts exhibited significantly increased binding of NF-κB p65 to the *Egr-1* promoter, suggesting that this treatment enhances NF-κB-mediated transcriptional regulation of Egr-1, potentially contributing to altered *Egr-1* gene expression under these conditions.

### 4-PBA induced the acetylation of histone H3 but not NF-κB p65

To determine whether 4-PBA treatment directly affects histone acetylation, we examined the acetylation status of histone H3 at residues H3K9, H3K14, H3K18, H3K27, and H3K56 during myogenesis by immunoblotting. Histone H3 acetylation markedly increased at the early stage of differentiation following inhibitor treatment, particularly at H3K18 and H3K27 (Figure 4A and 4B). These results suggest that HDAC inhibition by 4-PBA alters the acetylation status of specific histone lysine residues.

**Figure 4.**
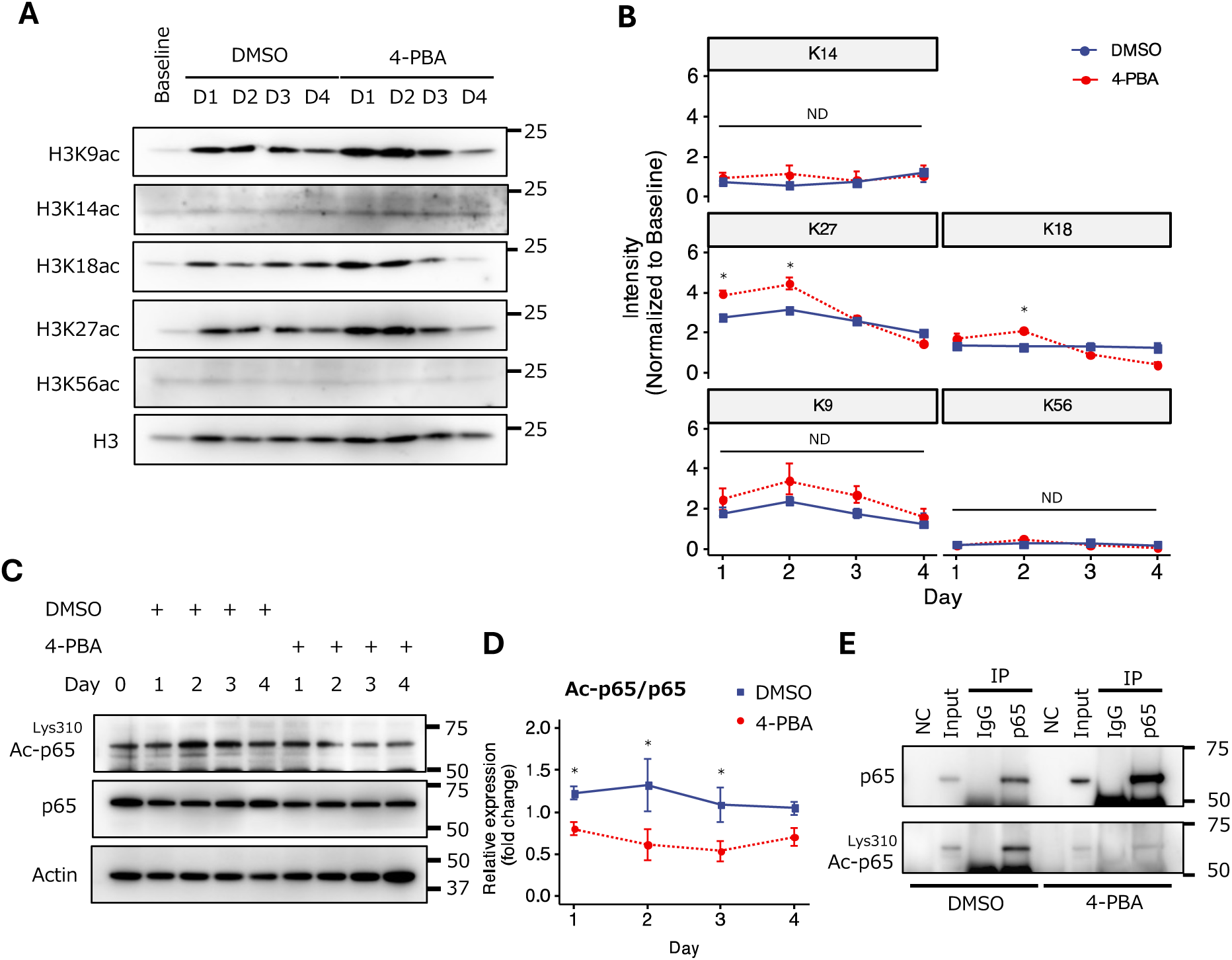
4-PBA Induces Histone H3K27/18 Acetylation but Does Not Promote p65 Acetylation. (A) Representative immunoblots showing acetylated histone H3 (H3K9/14/18/27/56ac) in C2C12 cells treated with 1 mM 4-PBA or DMSO for the indicated time points. Total histone H3 was used as a loading control. (B) Quantification of histone H3K27/18 acetylation shown in (A), normalized to total histone H3. Data represent three independent experiments. (C) Representative immunoblots showing total p65 and acetylated p65 (Lys310) in C2C12 cells treated with 1 mM 4-PBA or DMSO for the indicated time points. β-actin (Actin) was used as a loading control. (D) Quantification of acetylated p65 (Lys310) levels shown in (C), normalized to Actin. Data represent three independent experiments. (E) Immunoprecipitation of p65 from C2C12 cell lysates followed by immunoblot analysis with antibodies against total p65 or acetylated p65 (Lys310). Input lysates are shown as controls. Data in (B) and (D) are presented as mean ± S.E.M. and analyzed using an unpaired two-tailed Student’s t test (B, D). \**P* < 0.05; ns, not significant.

Other groups have reported that 4-PBA directly interacts with NF-κB p65 and regulates its acetylation^32^. We therefore evaluated the acetylation of NF-κB p65 at Lys310 during C2C12 myoblast differentiation by immunoblotting. The acetylation of NF-κB p65 at Lys310 increased during myoblast differentiation; however, inhibitor treatment reduced p65 acetylation (Figure 4C and 4D). Consistently, coimmunoprecipitation using an antibody against p65 in C2C12 myotubes revealed that inhibitor treatment decreased NF-κB p65 acetylation (Figure 4E). These findings indicate that despite its strong effects on histone acetylation, 4-PBA does not increase and instead reduces NF-κB p65 acetylation in C2C12 cells.

### 4-PBA-induced Hdac5 expression negatively regulates Egr-1 expression during myogenesis

4-PBA suppresses Hdac5 protein expression, leading to the hyperacetylation of histone H3 in C2C12 myotubes^33^. However, it is still unclear which class of HDACs is the primary target of 4-PBA. To investigate which classes of HDACs are predominantly downregulated by 4-PBA treatment during myogenesis, we first evaluated the gene expression of all the HDACs in C2C12 myotubes treated with 1 mM 4-PBA compared with those treated with DMSO. We observed significantly lower Hdac5 gene expression in C2C12 myotubes than in DMSO control myotubes (Figure 5A and Data S3). Hdac5 expression increases during the early stages of differentiation, confirming that Hdac5 expression is affected by the formation of immature myotubes (Figure 5B and 5C). In contrast, 4-PBA treatment reduced Hdac5 expression during myogenesis, which is consistent with the RNA-seq data. Together, these results suggest that Hdac5 is a key epigenetic regulator during early myogenesis and that 4-PBA treatment suppresses its expression, potentially leading to altered histone acetylation and dysregulated transcriptional programs during muscle differentiation.

**Figure 5.**
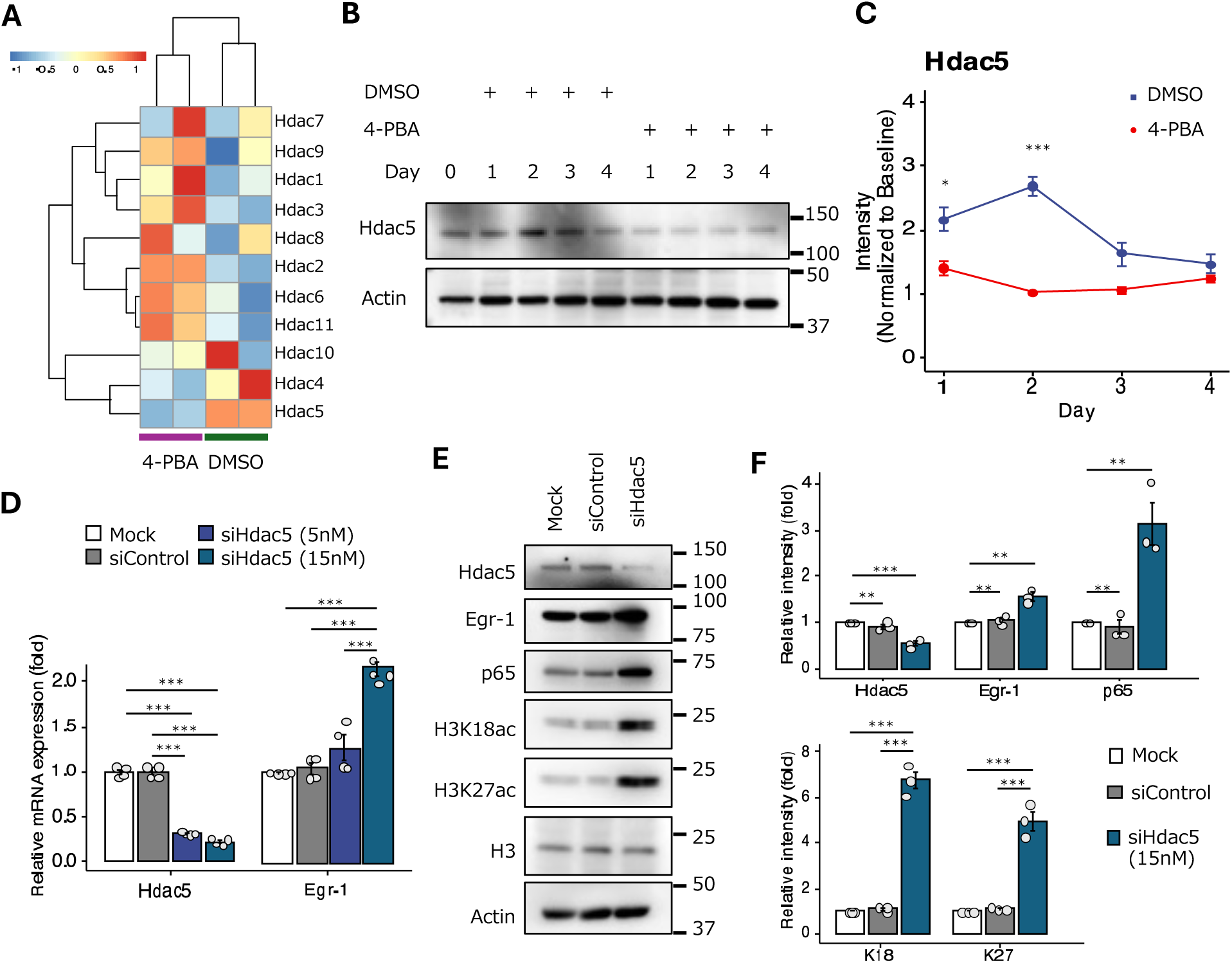
4-PBA Regulates Egr-1 Expression through Suppression of HDAC5 in C2C12 Cells. (A) Heatmap showing RNA-seq–based expression profiles of histone deacetylase (HDAC) genes in C2C12 cells treated with 1 mM 4-PBA or DMSO. (B) Representative immunoblots showing HDAC5 protein levels in C2C12 cells treated with 1 mM 4-PBA or DMSO for the indicated time points. β-actin (Actin) was used as a loading control. (C) Quantification of HDAC5 protein levels shown in (B), normalized to Actin. Data represent three independent experiments. (D) Relative mRNA expression of *Hdac5* and *Egr-1* in C2C12 cells transfected with siHdac5 (5 or 15 nM), siControl, or mock control, as measured by qPCR. Expression levels were normalized to *β-actin* (*Actin*). (E) Representative immunoblots showing HDAC5, Egr-1, p65, H3K18ac, and H3K27ac protein levels in C2C12 cells treated with 1 mM 4-PBA or DMSO for the indicated time points. Actin or histone H3 was used as the loading control, as indicated. (F) Quantification of protein levels shown in (E). Band intensities were normalized to Actin for HDAC5, Egr-1, and p65, and to histone H3 for H3K18ac and H3K27ac. Data represent three independent experiments. Data in (C), (D) and (F) are presented as mean ± S.E.M. and analyzed using an unpaired two-tailed Student’s t test (C) and one-way ANOVA with Tukey’s post hoc test (D, F). \**P* < 0.05, \*\**P* < 0.01, \*\*\**P* < 0.001.

We next confirmed that Hdac5 is directly related to Egr-1 expression. Depletion of *Hdac5* mRNA expression significantly increased the *Egr-1* mRNA level in 4-PBA-treated C2C12 cells (Figure 5D), suggesting that Hdac5 negatively regulates *Egr-1* expression. We next performed immunoblotting analysis to examine the protein levels of Egr-1 and NF-κB p65, as well as histone H3 acetylation at Lys18 and Lys27, which are known to be upregulated by 4-PBA treatment. The results revealed that the protein expression of Egr-1 and NF-κB p65, along with the acetylation of histone H3 at Lys18 and Lys27 (Figure 5E and 5F), was significantly greater in *Hdac5*-knockdown cells than in control cells. These results suggest that HDAC5 knockdown in C2C12 cells enhances histone H3 acetylation and activates NF-κB signaling and Egr-1 expression, indicating that the mechanism is similar to that of 4-PBA.

### 4-PBA regulated ECM organization-related genes via Egr-1

To investigate the mechanism through which Egr-1 affects myogenic differentiation, the gene expression profiles of C2C12 cells with or without 4-PBA were determined via RNA-Seq. GO enrichment analysis revealed that the significantly enriched functional categories among the upregulated genes included positive regulation of external encapsulating structure organization, extracellular matrix organization, extracellular structure organization and collagen aggregation (Figure 6A and Data S4). Furthermore, network analysis using GO enrichment revealed genes significantly involved in extracellular matrix (ECM) structural constituent, integrin binding and cell adhesion molecule binding (Figure 6B). In particular, the expression of collagen-related genes, which encode structural components of the extracellular matrix, increased during myoblast differentiation (Fig. S3A and Data S5), suggesting that ECM-associated genes play important roles in muscle differentiation.

**Figure 6.**
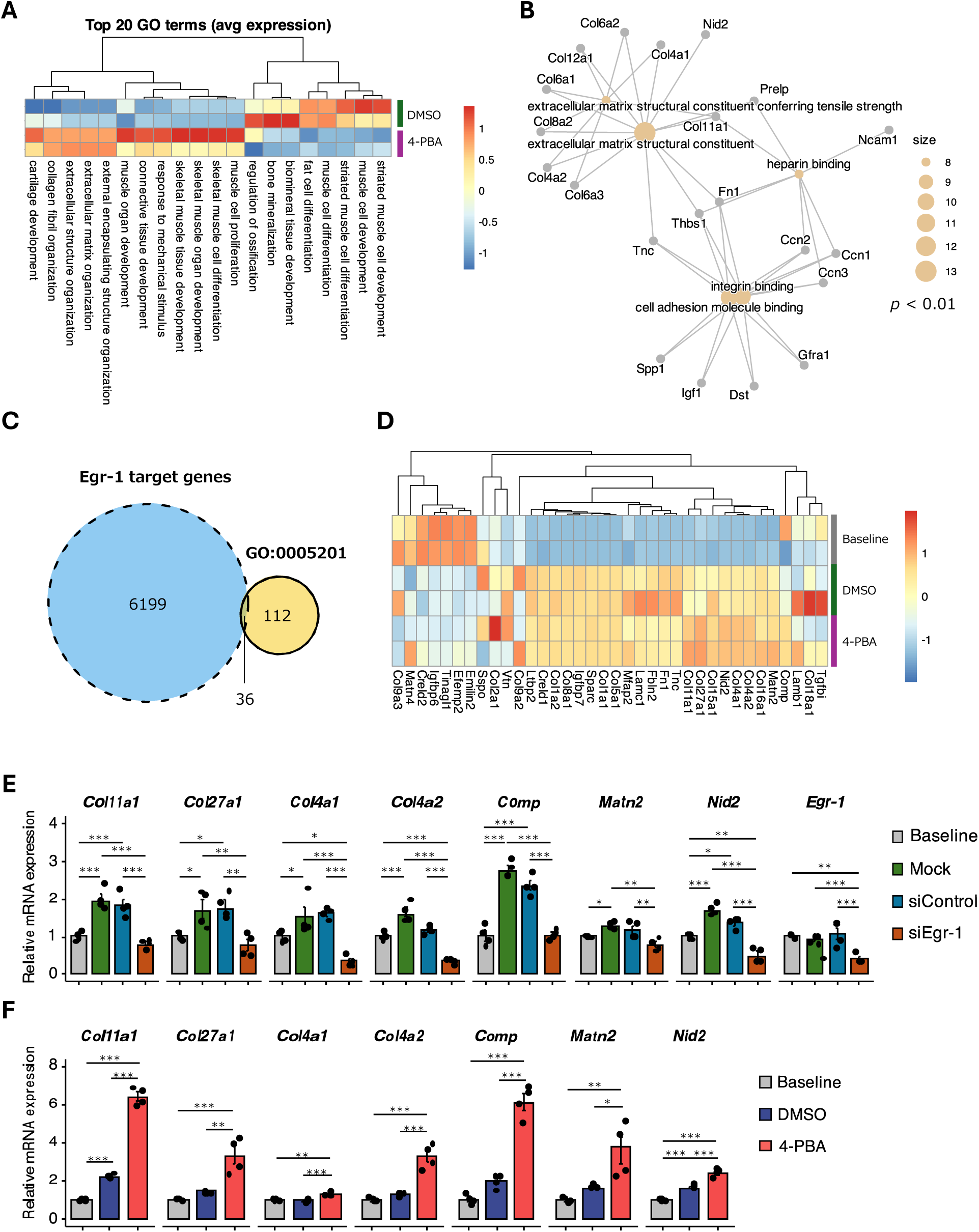
Egr-1 Mediates 4-PBA–Dependent Transcriptional Regulation of Extracellular Matrix Organization Genes. (A) Heatmap of Gene Ontology (GO) hallmark gene sets with the top 20 lowest p values identified by parametric gene set enrichment analysis (GSEA) from RNA-seq data of C2C12 cells treated with 1 mM 4-PBA or DMSO. (B) Network visualization illustrating the functional connectivity between differentially expressed genes (DEGs) and enriched biological processes, highlighting extracellular matrix (ECM) organization–related pathways regulated by 4-PBA. Node size corresponds to the number of genes within each enriched pathway. (C) Venn diagram showing overlap between genes associated with the ECM structural constituent GO term (GO:0005201) and predicted Egr-1 target genes obtained from the ENCODE Transcription Factor Targets Dataset. (D) Heatmap showing expression patterns of Egr-1 target genes within the ECM structural constituent gene set, demonstrating coordinated regulation downstream of 4-PBA treatment. (E) Relative mRNA expression of selected ECM-related genes in C2C12 cells transfected with siEgr-1 (15 nM), siControl, or mock control, showing Egr-1–dependent transcriptional regulation. Expression levels were normalized to β-actin (Actin). (F) Relative mRNA expression of selected ECM-related genes in C2C12 cells treated with 1 mM 4-PBA or DMSO under differentiation conditions. Baseline indicates expression levels prior to differentiation induction. Data in (E) and (F) are presented as mean ± S.E.M. and analyzed using a one-way ANOVA with Tukey’s post hoc test. \**P* < 0.05, \*\**P* < 0.01, \*\*\**P* < 0.001.

We next identified the ECM-related genes regulated by Egr-1 expression that were upregulated by 4-PBA treatment in C2C12 cells. A Venn diagram was constructed to examine the intersection between Egr-1 target genes based on the ENCODE Transcription Factor Targets Dataset^34^ and ECM structural components (GO: 0005201), indicating that a total of 36 common genes were identified (Figure 6C). On the other hand, we found that there were no genes common between Egr-1 target genes and categories for the adhesion molecule binding (GO:0050839), integrin binding (GO:0005178), and heparin binding (GO:0008201) that the network analysis using GO enrichment reveled (Figure S3B). To identify the genes linking 4-PBA to the upregulation of ECM via Egr-1 expression, we displayed the main representative up-regulated genes on a heatmap, and the expression of Egr-1 target genes by 4-PBA increased gradually from left to right (Figure 6D and Data S6). Changes in gene expression following Egr-1 knockdown were confirmed using qRT-PCR.

Several extracellular matrix (ECM)-related genes, including collagen type XI alpha 1 (Col11a1), collagen type XXVII alpha 1 (Col27a1), collagen type IV alpha 1 (Col4a1), and collagen type IV alpha 2 (Col4a2), as well as genes involved in the production of collagen, such as cartilage oligomeric matrix protein (Comp), and myogenesis-related genes, including matrilin-2 (Matn2) and nidogen-2 (Nid2), a crucial BM protein in muscle, were among the most significantly downregulated in Egr-1-transfected C2C12 myoblasts compared with those in siControl-transfected cells or at baseline levels (Figure 6E). Moreover, the expression of these genes was significantly greater in C2C12 myoblasts treated with the inhibitor than in those treated with DMSO (Figure 6F). Similar downregulation of these genes was also observed in siHdac5-treated cells (Figure S3C). Overall, these results suggest that 4-PBA promotes ECM remodeling and the expression of muscle-related genes and that Egr-1 is involved in regulating the expression of these genes.

## Discussion

Although significant progress has been made in understanding muscle differentiation, many of the underlying mechanisms remain unresolved. Here, we report that 4-PBA regulates myoblast differentiation, cell proliferation, and ECM remodeling via epigenetic and transcriptional mechanisms, advancing our understanding of muscle pathology and facilitating the development of novel therapeutic strategies for muscle-related diseases such as sarcopenia and muscular dystrophy.

We found that 4-PBA acts as an epigenetic regulator during myoblast differentiation. Transcriptomic screening and immunoblotting analysis revealed that histone H3 acetylation (H3K18ac and H3K27ac) increased during early differentiation, supporting the role of 4-PBA as an effective inhibitor of Hdac5 in C2C12 myoblasts. We demonstrate that compared with many HDAC inhibitors that reinforce muscle-specific transcription, 4-PBA inhibits myoblast differentiation and the expression of the myogenic-related genes *Myod* and *Dysf* to promote muscle differentiation. Importantly, 4-PBA selectively induced the expression of NF-κB p65 via inhibition of Hdac5 expression. NF-κB plays complex roles in skeletal muscle differentiation through opposing signaling pathways^35^. The canonical NF-κB signaling pathway, via RelA/p65, IKKβ, and IKKγ, and is a key transcription factor that generally promotes muscle cell proliferation, especially in skeletal muscle stem cells (MuSCs) after chronic injury^36–38^. We propose that 4-PBA promotes cell proliferation by activating the NF-κB signaling pathway, thereby contributing to myoblast expansion during early myogenesis following injury.

We identified HDAC5 as a key epigenetic regulator during early myogenesis and suggest that its suppression by 4-PBA contributes to altered histone acetylation and transcriptional dysregulation. We also suggest that 4-PBA is a target for the acetylation of histone H3 but not NF-κB p65 directly through HDAC5 inhibition. In fact, fasting or several types of muscular dystrophy increase p65 acetylation at Lys-310, activating NF-κB and contributing to muscle atrophy^39^. Acetylated NF-κB p65 also promotes the expression of inflammatory genes, a hallmark of muscle wasting^39,40^. In acute muscle injury, inflammation is necessary but must be precisely resolved to avoid chronic damage^41^. Notably, long-term 4-PBA treatment during myogenesis reduces the phosphorylation of NF-κB p65 gradually as myotubes mature, which is activated in the early stage (Figure 3A and B). These findings suggest that myogenesis-related factors may contribute to a negative feedback loop that controls the accumulation of NF-κB p65 signaling. This process contributes to the suppression of excessive activation of the NF-κB signaling pathway. 4-PBA can act downstream of many signaling cascades, allowing regeneration to proceed without broadly activating NF-κB-driven inflammatory pathways that can exacerbate tissue damage.

We identified Egr-1 expression as a key factor that alternates between cell proliferation and differentiation in myoblasts, and we functionally characterized the regulation of Egr-1 expression by NF-κB p65. We found that 4-PBA induces Egr-1-regulated extracellular matrix (ECM)- and collagen-related gene expression. Collagen provides structural stability to muscle fibers and the surrounding connective tissues, thereby contributing to myoblast differentiation, fusion, and proper structural organization^42^. In most muscular dystrophies, the accumulation of collagen, particularly type I, type III and type IV collagens, is markedly increased^43–46^. Excessive collagen deposition within the ECM, which is produced mainly by fibroblasts, progressively leads to fibrosis and impaired muscle function. In contrast, specific collagen types—primarily nonfibrillar collagens—serve as essential components of the MuSC microenvironment and regulate their behavior. In our study, we detected increased expression of Col27a1, Col4a1, and Col4a2, which are known ECM components that support normal muscle development, regeneration, and resistance to mechanical stress. Notably, Col27a1 is expressed in the MuSC niche, suggesting a potential role in regulating the local microenvironment^2^. Type IV collagen is a key component of the basement membrane (BM), which surrounds satellite cells together with laminin and interacts with surface α7/β1 integrins^47,48^. 4-PBA induces the expression of genes involved in collagen production, namely, Matn2 and Nid2. Matn2 functions as a key regulator of myogenic program initiation by modulating BMP7 and integrin α5 signaling, inducing the expression of the Cdk inhibitor p21, and activating myogenic genes, including Nfix, MyoD, and Myog^52^. Nid2 has been detected in conditioned media from differentiating human myoblasts by mass spectrometry, which suggests that it is secreted as a soluble protein during myogenesis^49,50^. Nid2 is also a component of the muscle basal lamina and plays an important role in neuromuscular junction development^51,52^. Collectively, these results indicate that 4-PBA promotes ECM gene expression via the NF-κB p65 - Egr-1 signaling axis, thereby facilitating the expansion of undifferentiated muscle cells and generating replacement cells capable of responding to injury-associated cues. Moreover, in 4-PBA-treated skeletal muscle cells, these specialized ECMs may regulate signaling activities and maintain satellite cells in an undifferentiated state.

HDAC inhibitors such as trichostatin A (TSA), sodium butyrate (NaB), and valproic acid (VPA) have been shown to influence myoblast differentiation by increasing histone acetylation and activating muscle-specific gene expression in muscular atrophy and muscular dystrophy models^53–55^. Recently, givinostat, known as DUVYZAT, which is an HDAC inhibitor for HDAC1 and HDAC3, was approved for patients with Duchenne muscular dystrophy and has the potential to restore dystrophin expression, reduce inflammation, and preserve muscle integrity^18^. Previous studies have shown that 4-PBA helps to restore the membrane localization of the DYSF protein and repair the sarcolemma in a mouse model of dysferlinopathy^27^. Our data also indicate that 4-PBA controls cell proliferation and differentiation by targeting histone H3 without hyperactivation of inflammatory pathways, including the NF-κB signaling pathway. Collectively, these findings, together with our present results, suggest that 4-PBA plays an important role in myogenic differentiation after injury through the regulation of gene expression related to ECM remodeling and the alleviation of ER stress, potentially contributing to the prevention of muscle diseases. In summary, this work establishes a role for the HDAC inhibitor 4-PBA in maintaining muscle homeostasis and identifies Egr-1 expression and NF-κB p65 as key molecules for promoting myoblast proliferation. Future studies are warranted to determine such potential clinical applications.

## Limitation of the study

Despite the mechanistic insights gained in this study, several limitations should be acknowledged, along with potential strategies to address them in future work.

First, Although we identified HDAC5 inhibition as a primary mechanism underlying 4-PBA-mediated effects on histone acetylation, NF-κB signaling, and ECM gene expression, 4-PBA is a multifunctional compound with pleiotropic biological activities. Our findings indicate that chemical chaperone activity and ER stress alleviation may act in parallel with epigenetic regulation. Dissecting these mechanisms will require complementary genetic approaches, including ER stress-specific modulators.

Second, our analysis of NF-κB signaling focused primarily on early myogenesis, where transient activation appears to promote myoblast proliferation. However, prolonged or excessive NF-κB activation is associated with chronic inflammation and muscle wasting. Long-term in vivo studies are therefore needed to determine whether 4-PBA maintains a beneficial balance between regenerative signaling and inflammatory resolution during aging or chronic muscle disease.

Finally, although we observed increased expression of ECM- and collagen-related genes associated with a supportive satellite cell niche, we did not directly assess ECM architecture, biomechanical properties, or interactions with fibroblasts and immune cells. Future studies combining histological, biophysical, and single-cell approaches will help clarify whether 4-PBA-induced ECM remodeling favors regeneration without promoting fibrosis.

Together, addressing these limitations will be critical for translating 4-PBA-based strategies toward safe and effective therapies for muscle injury and degenerative diseases.

## Methods

### Cell culture

C2C12, a myoblast cell line, were purchased from the American Type Culture Collection. Cells were cultured in growth medium containing Dulbecco’s Modified Eagle Medium (DMEM; Genesee Scientific) containing 10 % fetal bovine serum (FBS, Avantor Seradigm), 100 units/ml penicillin, and 100 µg/ml streptomycin (Tharmo Fisher Scientific). Cell lines were cultured in a humidified atmosphere at 37 °C in 5 % CO2, and the culture medium was changed every 2 days. When cells reached subconfluence, they were rinsed in phosphate-buffered saline (PBS, pH 7.4, Genesee Scientific), trypsinized with 0.25 % trypsin-EDTA solution (Corning) for 5 min at 37°C, and passaged at ratios of 1:4 - 1:8.

To induce the differentiation and cell fusion, cells were washed with PBS, then changed to differentiation medium containing DMEM containing 5 % horse serum, and 100 units/ml penicillin, and 100 µg/ml streptomycin. The differentiation medium was changed every day.

### Small interfering RNA (siRNA)

For knockdown of Hdac5 or Egr-1, we used Silencer Select Mouse Hdac5 (Assay ID s67423) or Mouse Egr-1 (Assay ID s65378) purchased from Invitrogen. C2C12 myoblasts were seeded at a density of 1 × 10^5^ cells/well into a 6-well plate and incubated with siRNAs (5 or 15nM) and Lipofectamine RNAiMAX transfection reagent (Invitrogen) according to the manufacturer’s protocol. The control was Silencer Select Negative Control No. 1 siRNA. siRNA-transfected cells were incubated at 37°C in 5% CO2 until the assay. Representative data from three independent experiments.

### RNA sequence

C2C12 myoblasts were plated at 7.0×10^5^ cells in 60 mm dishes in growth medium (DMEM with 10% FBS and penicillin/streptomycin). Once confluent, media was changed to differentiation medium (DMEM with 2% HS and penicillin/streptomycin) with 1mM 4-PBA or DMSO as control and left to differentiate for 4 days. Differentiation medium was replenished daily.

We purified total RNA using an Maxwell RSC simply RNA Cell Kit (Promega, AS1390), followed by DNase treatment and assess the RNA purity using a Tapestation 4200 High Sensitivity RNA (Agilent Technologies, 5067-5579).

RNA-seq analysis was performed as previously described. We used The NEBNext Poly(A) mRNA Magnetic Isolation Module (New England BioLabs, E7490) and NEBNext Ultra II Directional RNA Library Prep Kit for Illumina (New England BioLabs, E7760) to generate cDNA libraries for Illumina sequencing. we used An Agilent 4200 TapeStation (Agilent, D1000) to evaluate the quality and concentration of the libraries. Confirmed libraries were mixed in equal molar amounts for clustering and sequencing on an Illumina NovaSeq 6000 DNA sequencer with a 50-bp, using NovaSeq 6000 S1 Reagent Kit V1.5 (Illumina). We used CLC Genomics Workbench 23.0.2 for data analysis. Trimmed reads were mapped to the *Mus musculus* (GRCm39) release-110 database and counted according to default settings using a CLC Genomics Workbench (Qiagen). mRNA abundance is expressed as the log2-transformed transcripts per million plus one. We use the R package seqinr to read the FASTA file.

### Immunoblotting

Cultured cells were denatured with SDS sample buffer (Nacalai Tesque) at 99 °C for 5 min. Lysates were loaded 10 µg protein/lane to run SDS-PAGE. The proteins were separated by SDS-PAGE and transferred to polyvinylidene fluoride (PVDF) membranes. The membranes were blocked with Block Ace (Bio-rad) and incubated at room temperature for 1 hour with primary antibodies (Table S1). After washing PBS containing 0.1% Tween 20, the membranes were incubated at room temperature for 1 hour with horseradish peroxidase (HRP)-conjugated anti-rabbit or anti-mouse IgG (CST) as secondary antibodies. Signals were developed using an Immobilon Western Chemiluminescent HRP Substrate (Cytiva) and examined using an Amasham imaging system (GE Healthcare). To analyze the intensity of bands, we used Fiji (ImageJ 1.53t).

### Co-immunoprecipitation (Co-IP)

The cell lysates were extracted from the treated C2C12 myotubes by use of RIPA lysis buffer, and then cultured overnight with specific antibodies against p65 and IgG (negative control) in constant speed at 4 °C. Following mixing with magnetic-beads (Cytiva), the antigen-antibody mixture was acquired. After washing thrice in IP lysis buffer, immunoblotting was conducted for the eluted proteins. See Table S1 for primer antibodies details.

### Immunofluorescence

The cultured cells were fixed in pre-child 4% parahormardehid at room tempreture for 10 min. After washing in PBS containing 0.1 % Triton X-100 for 5 min, cells were incubated with 3 % bovine serum albumin (BSA, Sigma) at room temperature for 60 min to block non-specific binding and then incubated with Anti-Myh1 antibody (Sigma-Aldrich, ZRB1214) as a primary antibody at room temperature for 1 hour. After washing with PBS, the cells and sections were incubated with an Alexa Fluor 488 Goat Anti-Rabbit IgG H&L (abcam, ab150077) as a secondary antibody at room temperature for 1 hour. the cells and sections were enclosed using ProLong Glass Antifade Mountant (Thermo Fisher Scientific) after staining the nuclei with 4’, 6-diamidino-2-phenylindole (DAPI) at room temperature for 5 min. The fluorescent images were obtained under a fluorescence microscope (Keyence).

### Cell proliferation assay

C2C12 myoblasts were seeded in 96-well plates at 5,000 cells/well with 1mM 4-PBA or DMSO. After incubated for 24, 48, and 72 hours, cell proliferation was measured using Cell Titer 96 AQueous Assay Kit (Promega) according to the manufacturer’s instructions.

### Quantitative real time Polymerase Chain Reaction (qPCR)

Total RNA was prepared using the ISOSPIN Cell & Tissue RNA (Nippon Gene) and was transcribed into cDNA using the ReverTra Ace qPCR RT Master Mix with gRNA remover (Toyobo). The qPCR was performed using PowerTrack SYBR Green Master Mix for qPCR (Thermo Fisher Scientific, A46109) with specific forward and reverse primers in a QuantStudio3 (Thermo Fisher Scientific). The relative gene expression was calculated with the ΔΔCt method^56^ (Livak and Schmittgen, 2001), using GAPDH or Actin for normalization. See Table S2 for qPCR primer details.

### Chromatin immunoprecipitation (ChIP) - qPCR

ChIP DNA was obtained using The SimpleChIP Enzymatic Chromatin IP Kit (Cell Signaling Technology 9003) according to the manufacturer’s instructions. Briefly, 4 × 10^6^ C2C12 cells were treated with 4% PFA for 15 min for cross-link, and then with ultrasonic for shearing DNA into 500-bp of fragments. The immunoprecipitation was implemented with 30 µL magnetic beads for 2 hours, following overnight incubation of anti-p65 or control IgG antibodies. The collected precipitated chromatin was subjected to qPCR. See Table S3 for ChIP-qPCR for primer details.

### Quantification and statistical analysis

Statistical analysis were performed using R software (v4.5.1). We used Student’s t-test to compare differences between two samples. We also used one-way ANOVA and Tukey-Kramer test for differences between the variances of the three groups. Values are presented as means ± S.E.M.

Visualized and analyses were performed using the following packages: clusterProfiler (version 4.16.0), KEGGREST (version 1.48.1), and GO.db (version 3.21.0), ggplot2 (version 4.0.0), ggpubr (version 0.6.1), pheatmap (version 1.0.13), VennDiagram (version 1.7.3), org.Mm.eg.db (version 3.21.0) from Bioconductor (Huber W Nat Methods).

Gene Ontology (GO) enrichment analysis was performed using GO annotations^57^. Biological process terms were tested for enrichment using a hypergeometric test, and p-values were adjusted for multiple testing using the Benjamini-Hochberg method. GO terms with an adjusted p-value < 0.05 were considered significant. KEGG pathway enrichment analysis was evaluated using a hypergeometric test with all expressed genes as the background gene set^58^. P-values were adjusted using the Benjamini-Hochberg method, and adjusted p-values < 0.05 were considered significant.

## Acknowledgments

We thank T. Ogata for helping in RNA-seq data analysis and A. Kobayashi for supporting general lab work. We thank Science Research Center, Insutitute of Life Science and Medicine for animal research, and the Yamaguchi University Center for Gene Research for using Flow cytometry, Imaging system, Microplate reader, and Real-time PCR system. This work was supported by Astellas Foundation for Research on Metabolic Disorders (K.T.), Uehara Memorial Foundation (K.T.), New Frontier Research Grant by Yamaguchi University (K.T., N.T.), Finding-Out & Crystallization of Subliminals (FOCS) project by Yamaguchi University of Medicine (K.T., N.T.). This work was supported in part by a Grant-in-Aid for Research Activity Start-up (21K21219), Early-Career Scientists (22K17826) from Japan Society of the Promotion of Science (JSPS) (N.T.), ACT-X (JPMJAX222B) from Japan Science and Technology Agency (JST) (N.T.), the Ube City Next-Generation Researchers Project (N.T.). This work was the result of using research equipment shared in MEXT Project for promoting public utilization of advanced research infrastructure (Program for supporting construction of core facilities), Grant Number JPMXS0440400023.

## Author contributions

K.T. conceived and designed the study. K.T. and N.T. performed experiments and did the data analysis and interpretation. K.T. wrote the manuscript and prepared the figures. K.T. and N.T. reviewed and edited the paper.

## Competing interests

The author N.T. is a founder and director of ADDVEMO Inc.. ADDVEMO Inc. was not involved in the study design, data collection, analysis, interpretation of data, the writing of the report, or the decision to submit the article for publication. The author N.T. receives research funding from Otsuka Electronics Co. for this study. The author K.T. declares no competing interests.

## Data availability

RNA-seq data were deposited to GEO (GEO: GSE315556). All data in this published article (and its Supplementary Information files) are available.

## Notes

### Summary of Updates

Author affiliations updated. Figure 2 in the previous version was not displayed correctly due to a PDF formatting issue. This version corrects the figure display. No scientific content has been changed.

