## Supplementary Files for "4-Phenylbutyric Acid Activates an NF-κB - Egr-1 Axis to Control Myoblast Proliferation and ECM Gene Expression Profiles"

Corresponding author:

Kana Tominaga, Ph.D.

Yamaguchi University School of Medicine

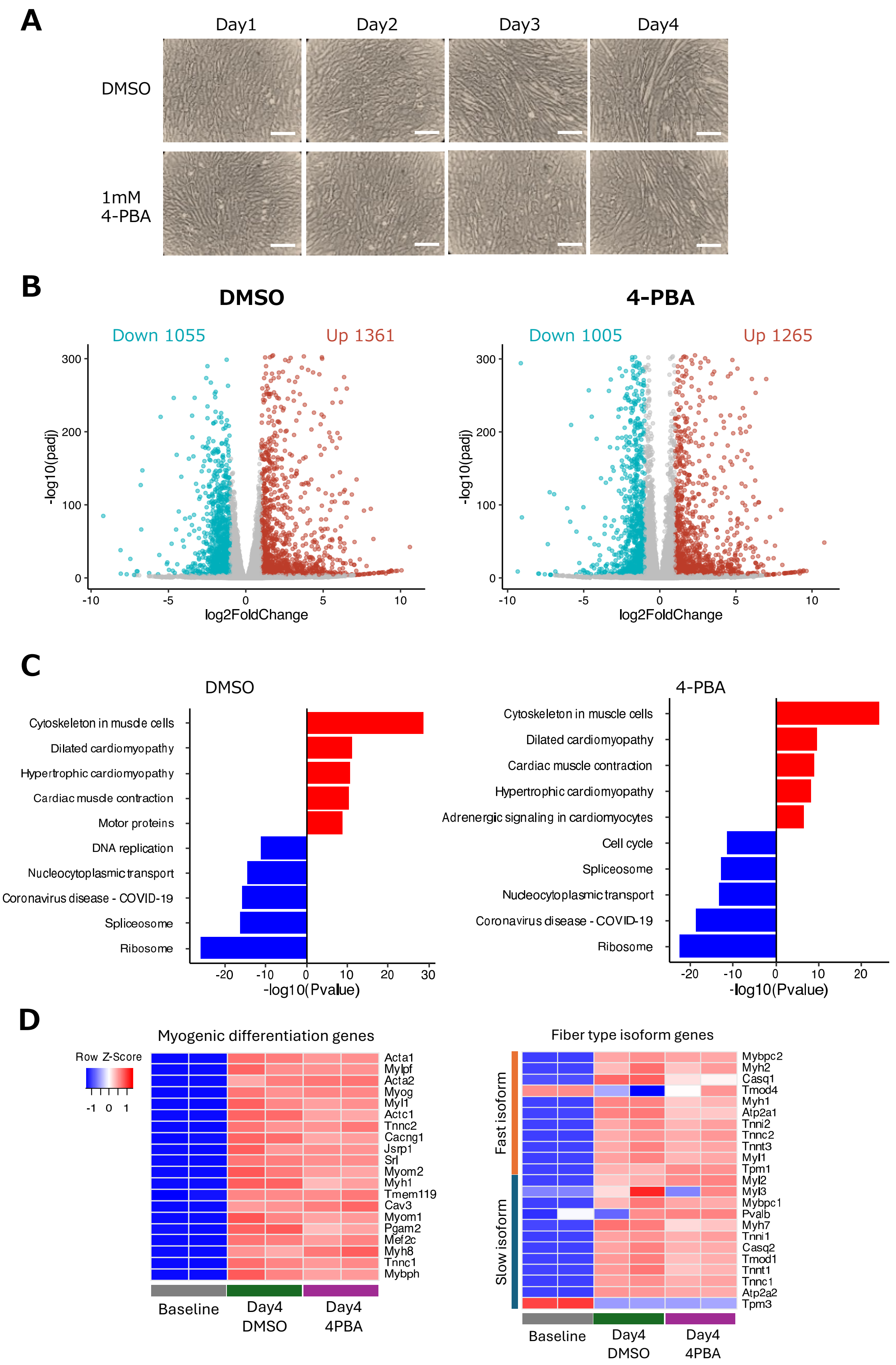

A
Day1
Day2
Day3
Day4
DMSO
1mM 4-PBA
B
DMSO
4-PBA
Down 1055
Up 1361
Down 1005
Up 1265
-log10(padj)
log2FoldChange
C
DMSO
4-PBA
Cytoskeleton in muscle cells
Dilated cardiomyopathy
Hypertrophic cardiomyopathy
Cardiac muscle contraction
Motor proteins
DNA replication
Nucleocytoplasmic transport
Coronavirus disease - COVID-19
Spliceosome
Ribosome
Adrenergic signaling in cardiomyocytes
Cell cycle
-log10(Pvalue)
D
Myogenic differentiation genes
Row Z-Score
Acta1
Mylpf
Acta2
Myog
Myl1
Actc1
Tnnc2
Cacng1
Jsrp1
Srl
Myom2
Myh1
Tmem119
Cav3
Myom1
Pgam2
Mef2c
Myh8
Tnnc1
Mybph
Baseline
Day4 DMSO
Day4 4PBA
Fiber type isoform genes
Fast isoform
Slow isoform
Mybpc2
Myh2
Casq1
Tmod4
Myh1
Atp2a1
Tnni2
Tnnc2
Tnnt3
Myl1
Tpm1
Myl2
Myl3
Mybpc1
Pvalb
Myh7
Tnni1
Casq2
Tmod1
Tnnt1
Tnnc1
Atp2a2
Tpm3

**Supplemental Figure S1. Effects of 4-Phenylbutyric Acid on Myogenic Differentiation and Gene Expression in C2C12 Cells, Related to Figure 1.**

(A) Representative phase-contrast images of C2C12 myoblasts treated with 1 mM 4-phenylbutyric acid (4-PBA) or DMSO under differentiation conditions. Scale bar, 400 µm.

(B) Volcano plot of differentially expressed genes identified by RNA-seq analysis, comparing 4-PBA- and DMSO-treated C2C12 cells before treatment (baseline) and at day 4 after myotube differentiation. Significantly upregulated genes are shown in red and downregulated are shown in blue. FC, fold change; P-adj, adjusted p value.

(C) KEGG pathway enrichment analysis of differentially expressed genes in 4-PBA- and DMSO-treated C2C12 myotubes, comparing with baseline and day 4 after myotube differentiation.

(D) Heatmap showing RNA-seq-based expression profiles of myogenic differentiation genes in C2C12 cells treated with 1 mM 4-PBA or DMSO at day 4 after myotube differentiation and at baseline (Day 0).

**A**

| sequence | start ~ end | strand |
|---|---|---|
| CTGGAATCCC | -162 ~ -171 | + |
| GGGAACTCCA | -1418 ~ -1427 | - |

**B**

**Mouse Gene Egr-1 (ENSMUST00000064795.6)**

TGGCCTGTTTGGGTCATGTGCCCATCCAAAAGTCATCACTGCAGGAAACAGAATGCTCATGGCTTACAAGGTCC

TATATGAATGGCCAGGCCTGGGTCACAGACCTACTCTTAGATTTTGGTGTGGAGTTCCCACAATCTCAGGAGTG

ACAGTAGCACCTCAGAAGAAATTGCTTACTGGGGAATATTCCAAAACTAACAGGTGCTGGGTACACAGAAGATA

CACACATGCAAATACCTCAGGCAGACTTGCACATCAGAGTCATGGGGGGAAATGAATATGATATTTAGAACTCT

ACTAGTTCTTAATACGACTTAGCCAGGGAGACAGAGACCTATGGGTTTATTTGAACCACTTTAGCACAATTACCA

GGCCATTACCCCTAAACTATCCCACACTCAAGTCACCAGCCAATCAGTTGTCCTGTCTGTGCTGGATCCACTCC

AGCCATGCCCTGCTTCTCCCCCCTCTTTCTCTTCCATGTGCCCCCTACCTACAGACCCTTCCCACCTCCCTACCT

CTGAAGCCCAGCCTACTTTTATTCTGCTCAGATCAGCCTCAGCCATCAGCAACTTTTTTTTTTTTAAGATTTATTTA

TTTTATGTATATGAGTACACTGTAGCTGTACAGATGGTTGTGAGCCATCATGTGGATTCTGGGAATTGAATTCAG

GACCTCTGCTCACTCCAGTTGGTCCTGCTCGCTCTGGCCCAAAGATTTATTTATTATTATTATCTAAGTACACTGT

AGCTGTCTTCAGACTCACCAGAAGAATGTGTCAGATCTCATTACAGATGGCTGTGAGCCACCATGTGGTTGTGG

GATTTGAACTCAGGATCTTTGGAAGAACAGTCAGTGCTCTTAACCGCTGAGCTATCTCTCCAGCTCCAATACTGT

CTTTTCAGTCCCCCACCGCGTGTCTCCAGTGCTTAGTAACGGGCAGTACGCGCTCTACATTTGGTAGTCACAGG

TCTGAACAAACAAACAATCCTTCCTCATGAAAACAGAGTTTACAACAAGAGCAGGTGAGACACAGGTTTGGTCA

GAGAATCCCACCAGAGTATCAAAATGGGCATGCTCTCACAGGAAGCACCTGGCCCTGCCTGTCCTGGCCAGGA

GGCTATGGAAAGGCACATTAATGGCTCCTAAATTAGTAACAACCAAAGAACTCAAACAGAATGGAACCCTGGTT

CATGTCTGGATTCCTTAAGCTGTTTTGGTTTTTTTAAATAAATAAAACGAAGGTGGAGGCGGATGGAGAGAGAGA

GAGAGAGAGAGAGAGAGAGAGAGAGAGAGAGAGAGAGAGCCGAAGAGTAACCTCCCTGTTAACCTTGACGTC

***P2_Forward***

CCCGATCTTCCTTCTTCTGTCTCTCAATCTCCTTCCACACAGGCACTCTCTGCTTTCTTTTTAAATAAAAAAATAAA

***P1_Forward*** ***P2_Reverse***

ATTAAAATAACAGCACCTTCCTCGTATTCAAAGTTGGAAACAAGAGCCTCCCATTCCTGGAATCCCTTCTCCCTT

***P1_Reverse*** ***P3_Forward***

TGGGTTGCTTCGGAGATAGGGCTTCACTGCTTGCGTCAGGGTCCCGGGAGACCAGCGGGATCTCTCTGCCATC

***P3_Reverse***

ACACCCCCGCCCCCTCCCCCCCCCCCCTGTTCCCTGCCCTTGGCCTGGCTCTGTGAAGGAAGTGTTACCCTG

AATTCTGGGCGCTTTGGCAGTGGCGGTTCCCTCGGGACTGCGGGGAAGGCCCAGGCCGCCG

**Supplemental Figure S2. Identification of a Putative NF-κB p65 Binding Site in the Murine *Egr-1* Promoter, Related to Figure 3.**

(A) Predicted NF-κB p65 binding site in the −2000 bp upstream region of the murine *Egr-1* promoter.

(B) Structure of the murine *Egr-1* promoter and primers (P1, P2 and P3). Predicted NF-κB p65 binding site is shown in blue. The T in red represents +1 site.

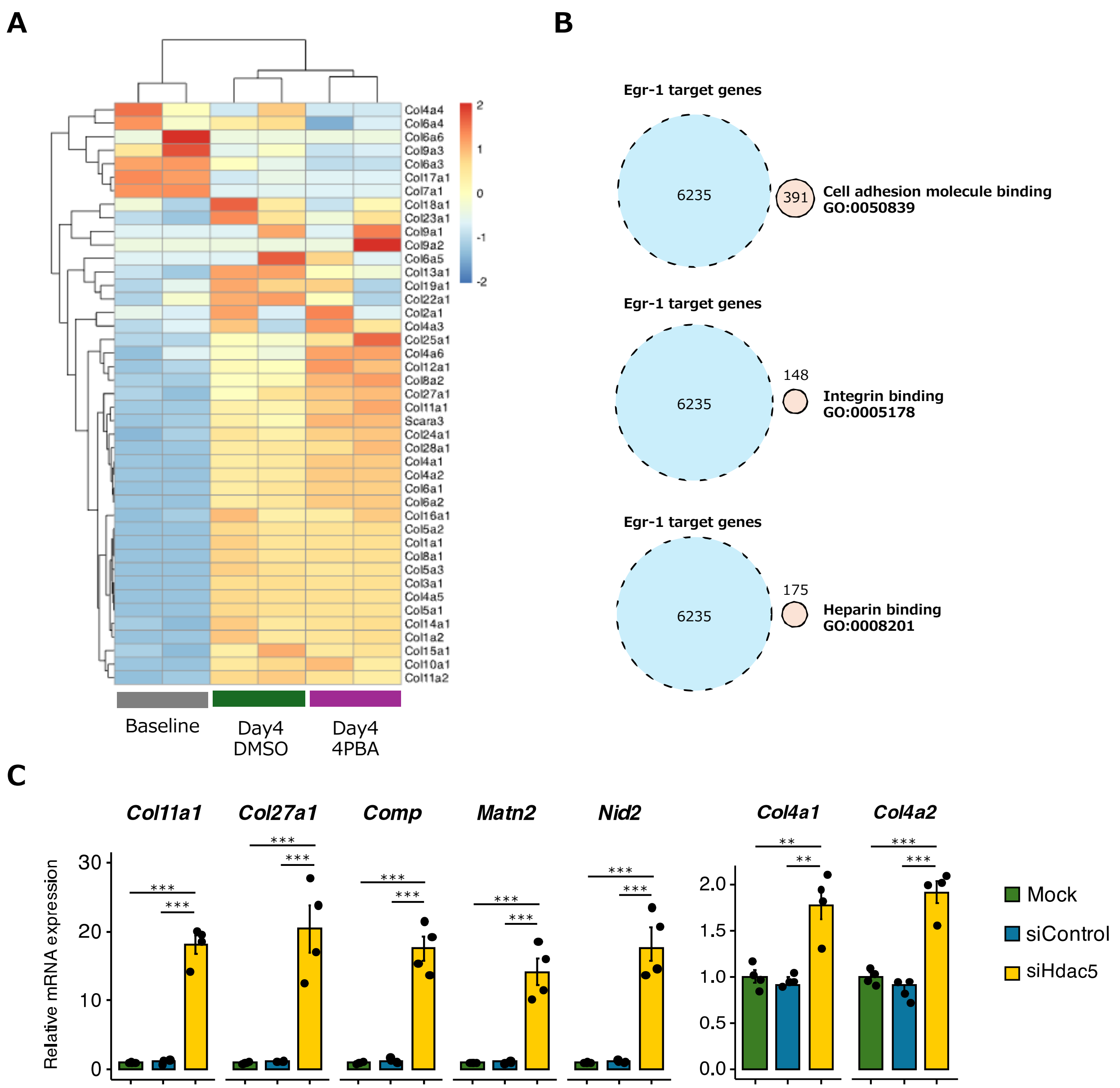


**Supplemental Figure S3. HDAC5–Egr-1 Axis Controls Expression of Collagen and Adhesion-Related Genes in C2C12 Cells, Related to Figure 6.**

(A) Heatmap showing RNA-seq-based expression profiles of collagen genes in C2C12 cells treated with 1 mM 4-PBA or DMSO at day 4 after myotube differentiation and at baseline (Day 0).

(B) Venn diagram showing overlap between genes associated with the the adhesion molecule binding (GO:0050839), integrin binding (GO:0005178), and heparin binding (GO:0008201) and predicted Egr-1 target genes obtained from the ENCODE Transcription Factor Targets Dataset.

(C) Relative mRNA expression of selected ECM-related genes in C2C12 cells transfected with siHdac5 (15 nM), siControl, or mock control, showing Egr-1–dependent transcriptional regulation. Expression levels were normalized to β-actin (Actin).

Data in (C) are presented as mean ± S.E.M. and analyzed using a one-way ANOVA with Tukey's post hoc test. *$P < 0.05$, **$P < 0.01$, ***$P < 0.001$.

**Table S1. List of antibodies in this study.**

| Antibody | Reference | Use and Dilution |
|---|---|---|
| Egr-1 | Proteintech,55117-1-AP | Western blotting (1:1000) |
| Fos | Proteintech, 66590-1-IG | Western blotting (1:500) |
| Myh1 | Sigma-Aldrich, ZRB1214 | Immunofluorescence (1:100) |
| Dysferlin | Leica Biosystems, NCL-Hamlet | Western blotting (1:1000) |
| Myogenin | BD Biosciences, 556358 | Western blotting (1:1000) |
| β-Actin | GeneTex, 14395-1-AP | Western blotting (1:10000) |
| Phospho-NF-κB p65 (Ser468) | Proteintech, 82335-1-RR | Western blotting (1:500) |
| Phospho-NF-kB p65 (Ser536) | Cell Signaling Technology, #3033 | Western blotting (1:1000) |
| Acetyl-NFkB p65 (Lys310) | Invitrogen, PA5-17264 | Western blotting (1:1000) |
| NF-κB p65 | Biolegend, 622602 | Western blotting (1:1000)<br>Immunoprecipitation (1:50) |
| Histone H3 (D1H2) | Cell Signaling Technology, #4499 | Western blotting (1:1000) |
| Acetyl-Histone H3 (Lys9) (C5B11) | Cell Signaling Technology, #9649 | Western blotting (1:1000) |
| Acetyl-Histone H3 (Lys14) (D4B9) | Cell Signaling Technology, #7627 | Western blotting (1:1000) |
| Acetyl-Histone H3 (Lys18) (D8Z5H) | Cell Signaling Technology, #13998 | Western blotting (1:1000) |
| Acetyl-Histone H3 (Lys27) (D5E4) | Cell Signaling Technology, #8173 | Western blotting (1:1000) |
| Acetyl-Histone H3 (Lys56) | Cell Signaling Technology, #4243 | Western blotting (1:1000) |
| HDAC5 (D1J7V) | Cell Signaling Technology, #20458 | Western blotting (1:1000) |

**Table S2. List of primers used for quantitative RT-PCR (qPCR) in this study.**

| Gene<br>NCBI | Forward (5' -> 3') | Reverse (5' -> 3') |
|---|---|---|
| Egr-1<br>NM_007913.5 | GTCCTTTTCTGACATCGCTCTGA | CGAGTCGTTTGGCTGGGATA |
| Fos<br>NM_010234.3 | CGAAGGGAACGGAATAAGATG | GCTGCCAAAATAAACTCCAG |
| Col4a1<br>NM_009931.2 | CTGGCACAAAAGGGACGAG | ACGTGGCCGAGAATTTCACC |
| Nid2<br>NM_008695.2 | CACCGAGGACAGTTTCCATT | CCAGTTACCAGGTGCTGGAT |
| Col11a1<br>NM_007729.3 | AGTTGGTCTGCAGTGGCAATTTCG | AGATCCCAGATCCACCGTTTCGTT |
| Col27a1<br>NM_025685.3 | CTCAAGGAGCCGTCAGATCG | AAGATCTCCCCTCCCTGGTC |
| Col4a2<br>NM_009932.5 | GACCGAGTGCGGTTCAAAG | CGCAGGGCACATCCAACTT |
| Matn2<br>NM_001358780.1 | GACGGACGGGCTCAGGAT | GATACCATTGGCCTTGGCTTTA |
| Comp<br>NM_016685.2 | GCGGTTCTATGAGGGTCCTG | GGGAGAAGCAGAAGACACCC |
| Hdac5<br>NM_001077696.2 | CCGGGAACCATCCTTGGAAA | CTTCACCTCCACTGCCACAG |

**Table S3. List of primers used for ChIP-qPCR in this study.**

| Name | Forward (5' -> 3') | Reverse (5' -> 3') |
|---|---|---|
| P1 | GTCCTTTTCTGACATCGCTCTGA | CGAGTCGTTTGGCTGGGATA |
| P2 | CGAAGGGAACGGAATAAGATG | GCTGCCAAAATAAACTCCAG |
| P3 | CTGGCACAAAAGGGACGAG | ACGTGGCCGAGAATTTCACC |
